# COVID-Align: Accurate online alignment of hCoV-19 genomes using a profile HMM

**DOI:** 10.1101/2020.05.25.114884

**Authors:** Frédéric Lemoine, Luc Blassel, Jakub Voznica, Olivier Gascuel

## Abstract

**Motivation:** The first cases of the COVID-19 pandemic emerged in December 2019. Until the end of February 2020, the number of available genomes was below 1,000, and their multiple alignment was easily achieved using standard approaches. Subsequently, the availability of genomes has grown dramatically. Moreover, some genomes are of low quality with sequencing/assembly errors, making accurate re-alignment of all genomes nearly impossible on a daily basis. A more efficient, yet accurate approach was clearly required to pursue all subsequent bioinformatics analyses of this crucial data.

**Results:** hCoV-19 genomes are highly conserved, with very few indels and no recombination. This makes the profile HMM approach particularly well suited to align new genomes, add them to an existing alignment and filter problematic ones. Using a core of ∼2,500 high quality genomes, we estimated a profile using HMMER, and implemented this profile in COVID-Align, a user-friendly interface to be used online or as standalone via Docker. The alignment of 1,000 genomes requires less than 20mn on our cluster. Moreover, COVID-Align provides summary statistics, which can be used to determine the sequencing quality and evolutionary novelty of input genomes (e.g. number of new mutations and indels).

**Availability:** https://covalign.pasteur.cloud, hub.docker.com/r/evolbioinfo/covid-align

**Supplementary information:** Supplementary information is available at *Bioinformatics* online.

## 1 Introduction

Since the emergence of the hCoV-19 virus (or SARS-CoV-2) responsible for the COVID-19 pandemic, unprecedented efforts are taking place across the world to sequence genomes of this virus and share the data. As of today (5/20/2020), the GISAID (Shu *et al*., 2017) provides access to more than 30,000 full genomes, and the NCBI and EBI more than 4,000 and 2,000, respectively. The first genomes were sequenced in China by the end of December 2019. Their number first increased slowly and then rapidly when the pandemic appeared on all continents. Submissions of several thousand sequences to GISAID in a single day has become common. Moreover, some genomes may be submitted incomplete, with sequencing and assembly errors. These characteristics pose major challenges to bioinformatics, notably that of multiple sequence alignment (MSA; Chatzou *et al.*, 2016), which is crucial for subsequent analyses (phylogeny, transmission clusters, mutation study, structure, etc.). To solve this difficulty, we use a profile HMM-based approach (Durbin *et al.*, 1998), which is the norm for HIV (www.hiv.lanl.gov), and is particularly well suited to hCoV-19, as its genome is highly conserved, without known recombination in human hosts (Xiaolu et al., 2020; De Maio et al., 2020). Using a profile, the addition of new data to an existing MSA requires linear computing times in the number of input genomes. Moreover, profile-based MSA proved to be very accurate (Earl *et al.*, 2014; Nute and Warnow, 2016). This approach is implemented in COVID-Align, which can be used thanks to a Web service and via Docker.

## 2 Methods

To estimate our profile HMM, we proceeded in several steps, in order to select an appropriate set of sequences and obtain a clean and reliable MSA to give as input to HMMER (www.hmmer.org):

- We downloaded all hCoV-19 genomes available on GISAID (April 24, 2020) and performed pairwise alignments using MAFFT (Katoh and Standley, 2013) of each of these genomes with the reference strain hCoV-19/Wuhan/WIV04/2019, sequenced in China December 30, 2019. This genome was found perfectly conserved not only in China, but also in Thailand, Japan, USA, UK, etc. and is considered as the origin of the virus (Li *et al.* 2020; www.gisaid.org).
- Then, using loose thresholds, we removed the genomes that were excessively divergent from the reference and had too many unknown (N) characters. We edited the remaining ones (e.g. removing the first gappy positions and the poly-A tail) and aligned them with MAFFT.
- The MSA so obtained was further filtered by removing the genomes having too many unique (i.e. not shared by any other genome) mutations and indels. We used more stringent thresholds than in the previous stage. This resulted in an MSA of 2,426 genomes, where the 12 first and 22 last positions of the reference genome were removed due poor alignment and low signal, but all other reference positions were preserved and showed high conservation. We used HMMER to estimate our profile from this curated MSA. All details and program options are available in Supplementary Information.

The resulting profile was implemented in a Nextflow (Di Tomaso *et al.* 2017) and Galaxy workflow combining hmmalign from HMMER to align the input genomes to the profile, GoAlign to format the input/output files (https://github.com/evolbioinfo/goalign), and Python to compute summary statistics. These statistics help users evaluate the sequencing quality and potential evolutionary novelties of input genomes; for example: number of unique mutations and indels, number of mutations compared to the reference genome… A user-friendly interface, implemented in GO (similar to Lemoine et al. 2019) allows users to launch their analyses without having to know how to use the Galaxy system. For advanced users, COVID-Align can be installed locally via Docker (https://www.docker.com).

## 3 Results

All results are given in a zipped file containing:

- The MSA of the input genomes plus the reference one that is displayed first, but cutting the first 12 and last 22 positions. With small datasets, this MSA can be visualized using MSAviewer (Fig. 1; Yachdav *et al.*, 2016).
- The hmmalign output in FASTA format, for each of the input genomes. This can be used to recover the insertions, deletions and match positions (to be reported to the reference genome).
- A CSV file with all statistics computed for each of the input genomes. Unique mutations and indels are possibly due to errors (sequencing, assembly etc.), while new ones (seen at least twice in submitted genomes, for the first time) likely correspond to evolutionary novelties (see Sup. Info. for details).
- A table in CSV format, summarizing the main average statistics and features of submitted genomes (Fig. 1).

**Figure 1.**
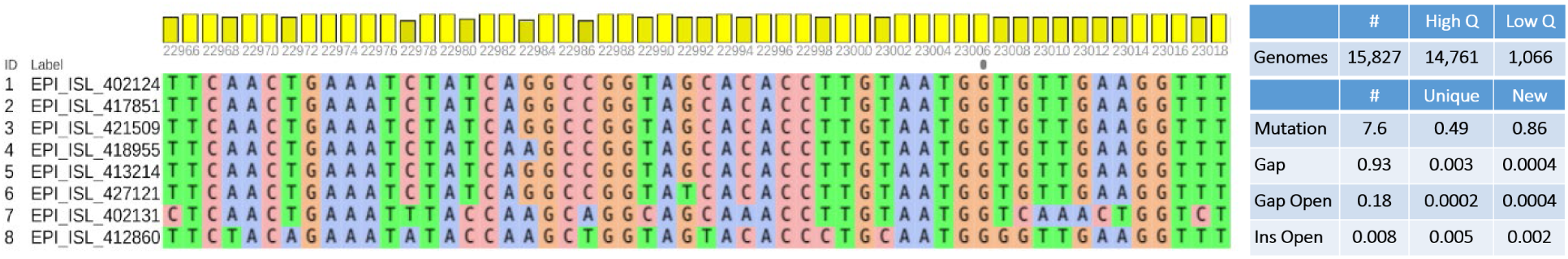
Visualization and Statistics Summary. Left: MSAviewer visualization of the Receptor Binding Domain (RBD) of the Spike gene, with reference genome (top), recently sequenced ones, and the Bat and Pangolin genomes (bottom). The site numbering corresponds to that of the reference, to be used to recover the ORFs and genes. In RBD region the Pangolin virus genome is closer to Human’s than is Bat’s, suggesting a possible recombination. On the opposite, Human viruses are highly conserved. Right: Statistics summary, displaying the number of High and Low Quality genomes, and the number of evolutionary events (mutations, gaps, gap openings, insertions, insertion openings). We distinguish the number of unique events (not seen yet and present only once in submitted genomes, possibly corresponding to errors) and the number of new events (seen at least twice, likely corresponding to evolutionary novelties). This table was filled with GISAID sequences deposited between April 25 and May 18, with unique and new statistics with respect to the database as of April 24 (Sup. Info.).

Our Web service processes 1,000 genomes in less than 20 minutes, thanks to parallelization that is easy to set up with profiles. Comparison with MAFFT-based GISAID MSA shows that our MSA: (1) can be used as is, while MAFFT’s cannot due to ∼10 000 highly gappy columns resulting from sequencing and assembly errors; (2) helps to detect and filter these errors; (3) is similar for most sequences to a properly trimmed version of MAFFT’s MSA, and more accurate for the few others (Sup. Info). Importantly, our profile and statistics will be regularly updated to account for user needs and the evolutionary novelties (mutations, indels…) of the emerging genomes to come.

## Acknowledgements

Sincere thanks to Amandine Perrin (Institut Pasteur) for her help, and the GISAID Team and all its Data Contributors for sharing their genome data.

## Funding

LB PhD Grant: PRAIRIE (ANR-19-P3IA-0001); JV PhD Grant: École Normale Supérieure Paris-Saclay and ED Frontières de l’Innovation en Recherche et Education, Programme Bettencourt; INCEPTION (PIA/ANR-16-CONV-0005).

## Conflict of Interest

none declared.

## SUPPLEMENTARY INFORMATION

### 1 Profile Estimation

To estimate our profile HMM, we proceeded in several steps, in order to select an appropriate set of sequences and obtain a clean and reliable MSA to give as input to HMMER (www.hmmer.org) *[we provide all details of this procedure below using bracketed, italic insertions in the main text]:*

We downloaded all hCoV-19 genomes available on GISAID (April 24, 2020; human host only) and performed pairwise alignments using MAFFT (Katoh and Standley, 2013) *[Options: mafft –add <seq>]* of each of these genomes with the reference strain hCoV-19/Wuhan/WIV04/2019 *[Genome ID: EPI_ISL_402124]*, sequenced in China December 30, 2019. This genome was found perfectly conserved not only in China, but also in Thailand, Japan and USA, and is considered as the origin of the virus (Li et al. 2020; www.gisaid.org). *[This genome serves as reference to curate daily submissions of new genomes on GISAID; several duplicates are available with 100% identity, but slightly shorter sequences resulting from different sequencing technology and submitter choices]*

Then, using loose thresholds we removed the genomes being excessively divergent from the reference and having too many unknown (N) characters *[a genome is removed if, compared to the reference, it has: >70 mutations, OR >15 internal indels (i.e. not situated at the sequence start and end), OR >20 start gaps, OR >20 end gaps, OR >50 ‘N’]*. We edited the remaining ones (e.g. removing the first gappy positions and the poly-A tail) *[positions 13 to 29,857 in the reference and pairwise aligned genomes are kept, positions 1-12 and 29,858-29,891 are eliminated]* and aligned them with MAFFT *[Options: mafft --thread 28 --auto <sequences>].*

The MSA so obtained was further filtered by removing the genomes having too many unique (i.e. not shared by any other genome) mutations and indels. We used more stringent thresholds than in the previous stage *[a genome is removed if in the second, global MSA it has: >3 unique mutations, OR >3 unique internal indels]*. This resulted in an MSA of 2,426 genomes, where the ∼40 first and last positions of the reference genome were removed due to poor alignment and low signal, but all other reference positions were preserved and showed high conservation *[>99.9% in average among all positions, but 48 variable positions with less than 90% conservation; average fraction of gaps per site = 0.05%, but 69 positions with more that 0.1% gaps]*. We used HMMER to estimate our profile from this curated MSA *[Options: hmmbuild -n covid19 covid19.hmm <alignment>]*.

### 2 Summary Statistics

For each of the input genomes, COVID-Align computes a series of summary statistics to help users analyze their data, remove problematic sequences, and detect those containing evolutionary novelties. As explained in the main text, we compute (among other statistics) the number of unique and new mutations/deletions/deletions. To achieve these computations, we regularly analyze all the data available on GISAID and count for every MSA position the number of A, C, G, T and gaps, and the number of times this position is followed by an insertion and the length of that insertion. When, a set of genomes is submitted, we compute the same quantities, which are used in combination with GISAID-based ones to obtain our summary statistics. Definitions are as follows:

- **A unique mutation/insertion/deletion** is present once and only once in the submitted sequences, but not in the GISAID sequences.
- **A new mutation/insertion/deletion** is either (1) not present in the GISAID sequences and seen at least twice in the submitted sequences, or (2) unique in the GISAID sequences and seen at least once in the submitted sequences. Importantly, <u>this does not apply to sequences already available on GISAID</u>, as these would be counted twice.

**The summary statistics returned for each of the submitted genomes** are as follows:

- **Length**: Length of unaligned sequence (not counting for starting/end gaps and unknown characters), to be compared to the length of the MSA (29,857 see above).
- **High_Quality**: Our quality index (Yes/No) based on the following rule: The sequence is deemed of high quality if it has : at most 8 unique mutations, at most 4 unique gap openings, at most 4 unique insertion openings, at most 40 mutations compared to the reference sequence, less than 5% N, less than 10% N + start gaps + end gaps.

#### MUTATIONS

- **Mut_Unique**: # Unique DNA mutations (see above definition for Unique/New).
- **Mut_New**: # New DNA mutations (does not apply to GISAID sequences).
- **Mut_Ref**: # DNA mutations compared to the reference genome (EPI_ISL_402124).
- **Mut_ORF:** # mutations occurring in ORFs.
- **Mut_Density**: Highest number of DNA mutations in a window of size 20 (to be used to detect poor quality genomes).
- **Mut_Unique_List:** List of unique mutations, as pairs of (position, nucleotide).
- **Mut_New_List:** List of new mutations (does not apply to GISAID sequences).
- **Mut_ORF_List:** List of mutations compared to the reference sequence, occurring in ORFs. Each mutation is represented as a triple of (position, mutated Nucleotide, name of ORF).

#### GAPS

- **Gap_Start**: # Gaps (i.e. deletions) at the beginning of the sequence (not counting those in the 12 first positions of the reference sequence).
- **Gap_End:** # Gaps at the end of the sequence (not counting those in the 22 last positions of the reference sequence).
- **Gap**: # Gaps in the core sequence (i.e. not counting start/end gaps).
- **Gap_Unique**: # Unique core gaps (see above definition for Unique/New).
- **Gap_New**: # New core gaps (does not apply to GISAID sequences).
- **Gap_Opening:** # Number of core gap openings.
- **Gap_Opening_Unique:** # Number of unique core gap openings.
- **Gap_Opening_New:** # Number of new core gap openings.
- **Gap_ORF:** # gaps occurring in ORFs.
- **Gap_Segment_Unique**: # Unique gap segments in the core sequence, having a unique set of starting position and length (see above definition for Unique/New).
- **Gap_Segment_New**: # New gap segments in the core sequence (does not apply to GISAID sequences).
- **Gap_Unique_List:** List of unique gap positions
- **Gap_New_List:** List of new gap positions
- **Gap_Opening_Unique_List:** List of unique opening gap positions
- **Gap_Opening_New_List:** List of new opening gap positions
- **Gap_Segment_List:** List of gap segments as pairs of (starting position, length), including gap segments at the start and end of the sequence.
- **Gap_ORF_List:** List of gaps occurring in ORFs, as pairs of (position, ORF name).

#### INSERTIONS

- **Insertion**: # non N insertions in the core sequence (i.e. not counting start/end insertions).
- **Insertion_Opening**: # Core insertion openings.
- **Insertion_Opening_Unique**: # Unique core insertion opening positions (see above definition for Unique/New).
- **Insertion_Opening_New**: # New core insertion opening positions (does not apply to GISAID sequences).
- **Insertion_ORF:** # Insertion opening in ORFs.
- **Insertion_Segment_Unique**: # Unique insertions segments in the core sequence, having a unique set of opening position and length (see above definition for Unique/New).
- **Insertion_Segment_New**: # New insertions segments in the core sequence (does not apply to GISAID sequences).
- **Insertion_Opening_Unique_List:** List of unique opening insertion positions.
- **Insertion_Opening_New_List:** List of new opening insertion positions.
- **Insertion_Segment_List:** List of insertion segments as pairs of (opening position, length).
- **Insertion_ORF_List:** List of insertions occurring in ORFs, as pairs of (opening position, ORF name).

#### NUCLEOTIDE CONTENTS

- **A:** # A in the whole sequence.
- **C:** # C
- **G:** #G
- **T:** # T
- **N:** #N
- **W:** #W
- **S:** #S
- **M:** #M
- **K:** #K
- **R:** #R
- **Y:** #Y
- **Ambiguous_Bases:** # ambiguous bases.

#### AVERAGE RESULTS

From these statistics, we compute average results for all submitted genomes (CSV format):

|  | # | High Q | Low Q |
| --- | --- | --- | --- |
| Genomes | Number of submitted genomes | Number of *High_Quality* genomes | Number of *Low_Quality* genomes |
|  | # | Unique | New |
| Mutations | Average *Mut_Ref* | Average *Mut_Unique* | Average *Mut_New* |
| Gap | Average *Gap* | Average *Gap_Unique* | Average *Gap_New* |
| Gap Open. | Average *Gap_Opening* | Average *Gap_Open_Unique* | Average *Gap_Opening_New* |
| Ins. Open. | Average *Insertion_Opening* | Average *Insertion_Opening_Unique* | Average *Insertion_Opening_New* |

For example, in the following Table (also provided in main text) we display the average statistics obtained for all available GISAID sequences from human host added between April 25 and May 18 2020, with “Unique” and “New” statistics based on all available GISAID sequences up to April 24.

|  | # | High Q | Low Q |
| --- | --- | --- | --- |
| Genomes | 15,827 | 14,761 | 1,066 |
|  | # | Unique | New |
| Mutations | 7.6 | 0.49 | 0.86 |
| Gap | 0.93 | 0.003 | 0.0004 |
| Gap Open. | 0.18 | 0.0002 | 0.0004 |
| Ins. Open. | 0.008 | 0.005 | 0.002 |

These results confirm that insertions are very rare. The number of shared insertion openings is 0.2% per genome, that is, 28 in total, with length of 1 to 3 nucleotides, corresponding to 11 sequences. Most of them are shared by 2 or 3 sequences only, and could be sequencing or assembly errors. Only one insertion of length 3 found in ORF 1ab is shared by 5 sequences from UK and Australia. This contrasts with deletions (gaps), which are much more frequent, with some long shared deletions, e.g. the 382-nt deletion found in over a dozen sequences from Singapore and Taiwan. When new sequences with confirmed insertions and deletions will be available from emerging genomes, they will be incorporated in the profile and the resulting MSA will closely account for these indels.

The “New” statistics shown in above table are based on all human sequences available on GISAID up to April 24. This is intended to illustrate the behavior of COVID-Align and the type of results the users should expect. But in real use, these statistics are based on a database that is regularly updated top account for the last evolutionary events observed among emerging genomes.

In above table, COVID-Align was used to align, assess quality and summarize features of newly submitted sequences sampled from human hosts. Nevertheless, COVID-Align can also provide high quality alignments of sequences sampled from various animal hosts, environment or cell cultures, or even sequences of more distant viral species from Coronaviridae family. Furthermore, as an HMM profile, it can be used to search for related sequences in a data pool.

### 3 Comparison with MAFFT-based GISAID MSA; trimming poor sequences

The genomes submitted to the GISAID and the other repositories may be incomplete with assembly and sequencing errors, and long stretches of unknown characters and gaps. These unusual characteristics, in addition to the number of genomes available, make multiple alignment difficult, despite the fact that these genomes are highly conserved up to now, after ∼6 months of evolution. The sequences need not only to be compared and aligned, but also to be trimmed. In that respect, a profile HMM approach is especially well suited.

The GISAID web site provides an MSA of all high-quality (<5% N) complete genomes (>29,000 length), without duplicates. This MSA is inferred using MAFFT Version 7 (Katoh and Standley, 2013) with Options: --thread −1 --nomemsave. We downloaded the MSA available on April 26, comprising 15,290 sequences. This MSA is much longer (38,570 sites) than the genome length (EPI_ISL_402124 = 29,891). This is due to the low quality of certain sequences. Even if these sequences were manually curated and assessed to be of high quality, they still contain a relatively large fraction of unknown characters (N), gaps (-) and assembly errors. Moreover, the length of N stretches seems to be approximate and poorly correlated to the real length of the corresponding sequences. This MSA thus contains a large number of columns containing ∼100% gaps, but for one or a few (mostly N) characters. Figure 1 provides an example where a large number of gaps is caused by an assembly error, plus a long stretch of unknown N characters. In some cases, likely due to the combination of poor sequences and progressive alignment strategy, MAFFT produces difficult to understand errors, as in Figure 2 where a portion of sequence with perfect match is shifted, resulting in a number of mismatches and gaps. Moreover, as noticed by several groups, the beginning and end of this MSA (and any other) are of particularly low quality, due to incomplete sequences, poly-A tails, etc. Consequently, this MSA cannot be used as is for most applications, e.g. to infer phylogenies.

**Figure 1:**
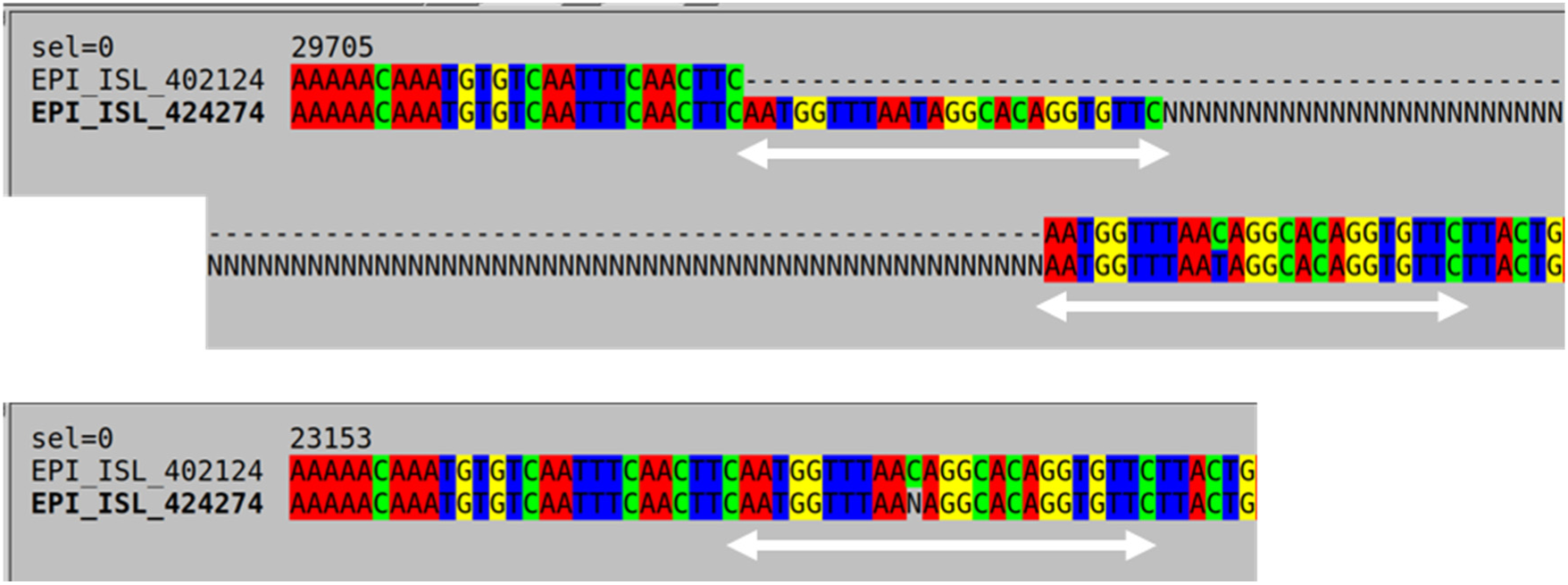
MAFFT (top, untrimmed) versus COVID-Align (bottom) MSA extracts with assembly error. EPI_ISL_402124 is the reference genome. Sequence EPI_ISL_424274 most likely contains an assembly error, as the segment AATGGTTTAATAGGCACAGGTGTTC is repeated twice. This, combined with a long stretch of N (unknown) characters, creates a large number of gaps in the reference sequence and in the whole MSA. COVID-Align detects this error as an insertion (unique among all GISAID sequences) and does not have any gap in this region.

**Figure 2:**
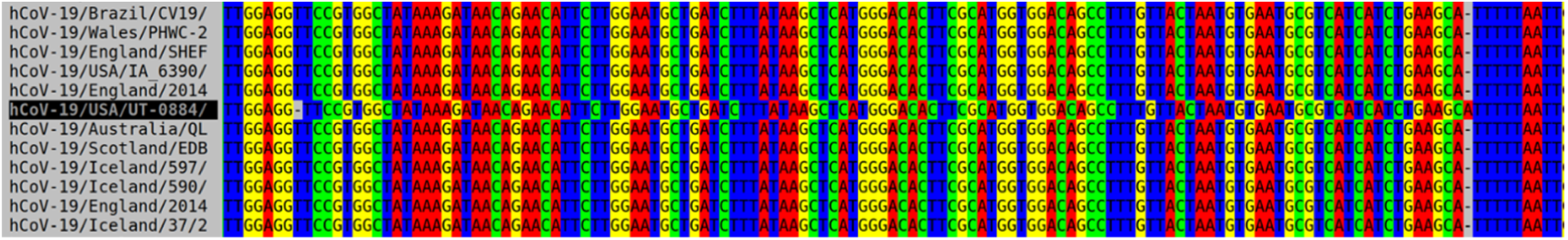
MAFFT (untrimmed) MSA with shift. A portion of the sequence is shifted, while in this region this sequence does not contain any insertions, gaps or N characters. In this region COVID-Align produces a perfect match, as expected.

On the opposite, COVID-Align MSA starts at position 13 in the reference genome, stops at position 29,869, and has a fixed length of 29,857. Assembly error in Figure 1 is trimmed, and the shifted region in Figure 2 is perfectly aligned. All sites in the MSA are highly conserved. Insertions are short and very rare, while some long deletions are found and confirmed as they are observed in several sequences of different origins (see above). Our profile will be updated regularly. If well-assessed insertions and deletions are found (as expected) in new emerging genomes, they will be added to the profile to reflect these important features of genome diversity.

To compare the two MSAs on the same basis, we trimmed MAFFT’s by removing all columns corresponding to gaps in the reference genome, as well as the first 12 and last 22 reference positions. Thus, both MSAs have the same length, refer to the same position in the reference genome and become similar, with 13,788 sequences having 100% identical alignment, and 1,499 sequences showing at least one mismatch (two different characters at the same position; N and gap characters are considered the same due to ambiguities and errors in the input sequences). Visual inspection shows that most differences between both MSAs are situated at the beginning and end of the sequences, due to N characters, poly-A tails, incompleteness of the sequences, etc. Thus, for each MSA we searched in each of the 1,499 differing sequences for the “real start” and “real end” of the aligned part of the given sequence, that is, the first and last windows of length 10 with at most 1 mismatch with the reference genome. When both MSAs indicated different start/end, we used the common part. Restricting the comparison to this common part, 1,417 sequences have identical alignment, and 75 show at least one mismatch. Moreover, the discarded parts (before the “real start” and after the “real end”) represent a very small fraction (∼0.2%) of the 1,499-sequence MSA, with ∼93% mismatch in average with the reference genome. On the opposite, the conserved part (∼99.8% of the MSA) has ∼0.5% mismatch in average with the reference genome. This confirms that the discarded start and end parts contain too many sequencing errors and uncertainties to be used in most analyses.

We compared both MSAs for the 75 genomes with core differences, using the number of substitutions with the reference genome (a substitution is a difference at the same position between two A, T, G, C characters; gaps are not considered as they are sometimes confounded with unknown N characters; moreover, after trimming of MAFFT’s both MSAs have the same length). Results are displayed in Figure 3. To summarize, 5 genomes are slightly better aligned by the trimmed version of the MAFFT MSA (with differences of at most 2 substitutions), and 70 are better aligned by COVID-Align (with differences up to 123 substitutions). For example, for the extreme sequence (ID EPI_ISL_419249), COVID-Align has 11 substitutions with the reference genome, while the MAFFT MSA has 134 substitutions. Figure 4 displays a portion of both MSAs with strong differences. While this sequence is evolutionary close to the reference genome, it will appear as one of the most distant using the trimmed MAFFT MSA. Even if the number of such sequences is relatively low, the presence of these alignment errors will profoundly perturb analyses.

**Figure 3:**
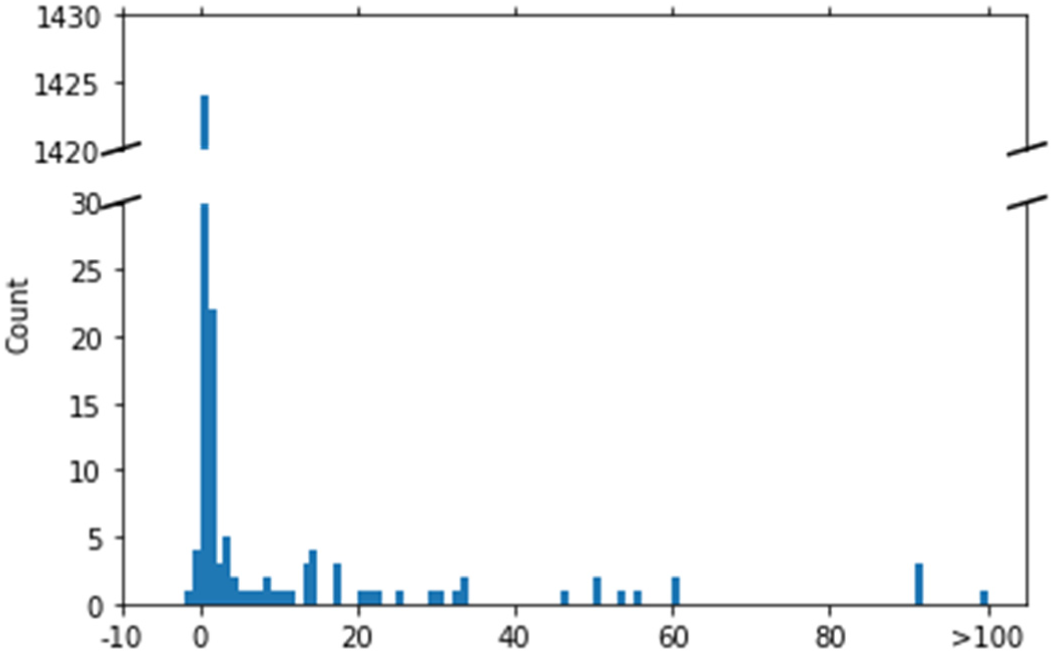
Difference in number of substitutions between COVID-Align and the trimmed MAFFT MSA.

**Figure 4:**
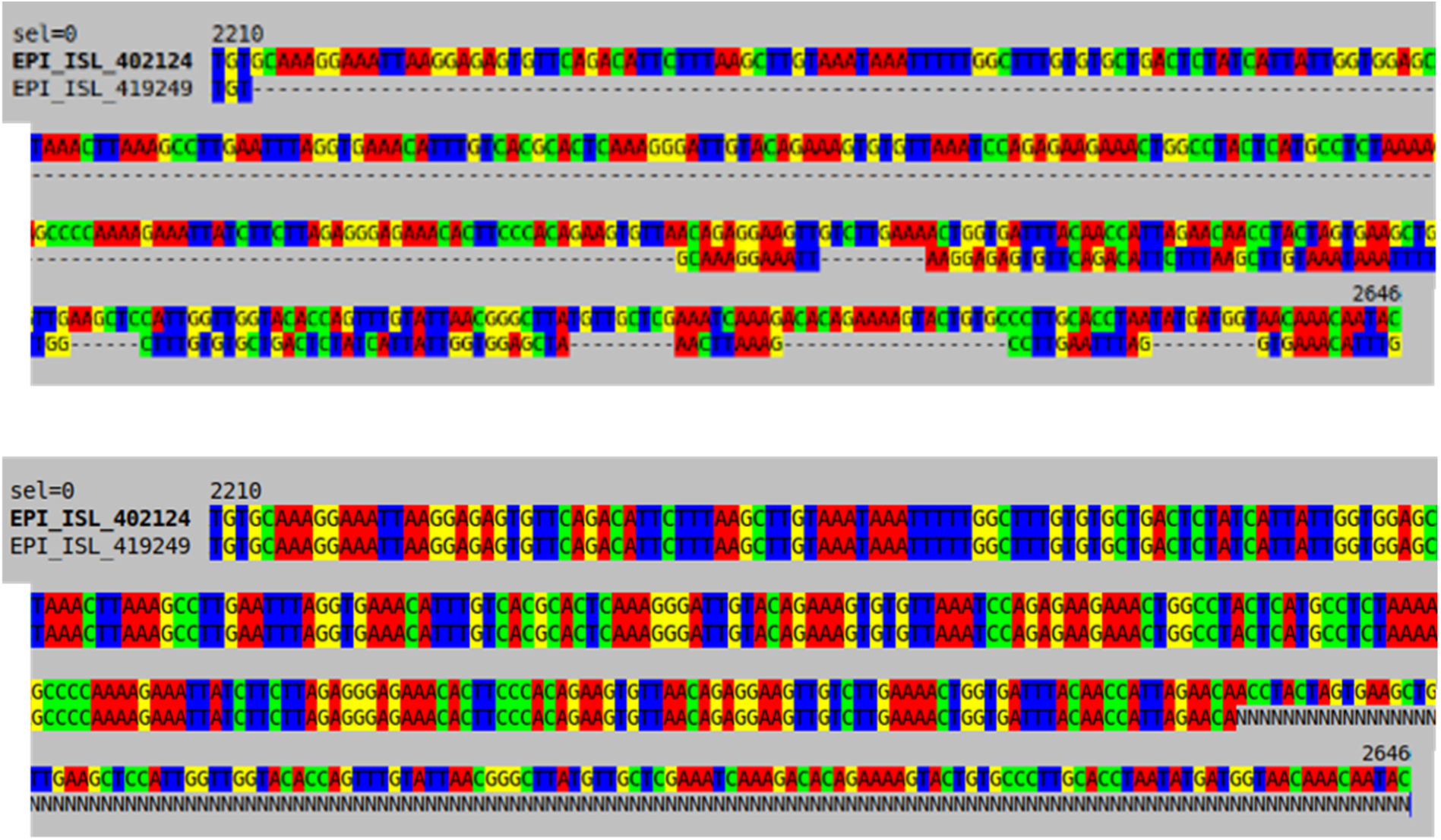
MAFFT (top, trimmed) versus COVID-Align (bottom) MSA extracts with sequence EPI_ISL_419249 and reference EPI_ISL_402124. While in this region EPI_ISL_419249 is very close to the reference, the MAFFT MSA introduces a number of gaps and substitutions.

To summarize, these results show the importance of trimming the MSAs obtained using MAFFT and any other standard aligner, and the accuracy of our profile HMM approach in both aligning the sequences and trimming the poorly sequenced or assembled regions, thus providing an MSA that is ready to use for further evolutionary and phylogenetic studies.

## Notes

### Competing Interest Statement

The authors have declared no competing interest.

